# Probing RNA-protein interactions in the early mouse embryo

**DOI:** 10.64898/2026.04.30.721999

**Authors:** Pablo Bora, Oliver J. Rando

## Abstract

The union of two germ cells to form a zygote, and subsequent early embryo development, are marked by radical remodeling of virtually every major class of biomolecules as the specialized germline states give way to the rapid and active growth that marks early development. In recent years, advances in ultra-low input genome-wide methods have enabled systematic analyses of mRNA abundance, and of chromatin organization, throughout early development in a variety of model systems. Here, we extend these efforts to the study of RNA binding protein (RBP) function in early mouse embryos, adapting REMORA ^1^ – based on fusing an RNA-editing enzyme to an RBP of interest – for use in early embryos. We benchmark our approach for several well-studied RBPs, successfully recovering expected features of their RNA cargos, and assayed the RNA cargos for 17 RBPs of interest for early gene regulation. Analysis of changes in mRNA metabolism following knockdowns of the RBPs surveyed here allowed us to identify direct regulatory functions for a subset of RBPs in the early mammalian embryo, including an unanticipated role for the RNA export adaptor Alyref in control of 3’ polyadenylation sites. Together, our data provide a proof of concept resource for systematically exploring RBP functions in mammalian embryogenesis.

## INTRODUCTION

Early preimplantation development in mammals is marked by dramatic and extensive molecular remodeling events to repackage the sperm genome, awaken the oocyte from arrest, and begin the complex choreography required to produce a fully-formed multicellular organism. Until recently, systematic molecular characterization of early mammalian embryos has been hampered by the limited numbers of oocytes or embryos that can be collected from one adult female (∼20-25 in mouse), and the small size of mammalian embryos, which has limited most analyses of the early mammalian embryo to microscopy-based readouts. That said, the past decade has seen remarkable technical advances in ultra-low input genomic approaches that have enabled detailed characterization of embryo transcription ^2–4^, chromatin accessibility ^5^, histone modification dynamics ^6–9^, and chromosome folding ^10–14^, providing spectacular insights into the processes by which the epigenetic states in germ cells are disassembled to allow establishment of the embryonic regulatory program, from embryonic genome activation to lineage allocation ^15^. More recently still, cutting-edge low-input proteomic technologies have been applied to the early embryo ^16,17^. Despite these advances, many regulatory systems remain inaccessible to current systematic analyses, as the limited material available precludes typical biochemical approaches.

Post-transcriptional gene regulation plays a unique role in early embryo development, as a store of deadenylated maternal transcripts are readenylated for translation ^18,19^, then subsequently degraded to make way for the new transcripts from the embryo’s genome, with each of these processes impacting specific subsets of transcripts in different ways. RNA-binding-proteins (RBPs) thus play central roles in the early embryo, modulating cell fate, zygotic genome activation (ZGA), and the morula to blastocyst transition, among other processes ^20–25^. Beyond RNA production and degradation, other regulatory aspects of RNA metabolism – from co-transcriptional splicing to alternative polyadenylation to covalent RNA modifications – are also likely to play key roles in early embryo development. However, systems-wide studies of RNA-RBP interactions in the early embryo have remained challenging in mammals, where limited input material has prevented application of CLIP or other biochemical assays of RBP binding to RNA targets. As a result, the functions for RBPs in shaping the co- and post-transcriptional RNA lifecycle in mammalian embryos remain largely underexplored.

Over the past decade, an alternative set of approaches to studying RBP-RNA interactions has been developed based on fusing RBPs to RNA base editors, allowing identification of RBP-bound RNAs based on edits that can be readily identified by RNA sequencing, thereby bypassing biochemical crosslinking and antibody pulldowns ^1,26–29^. The RNA editors used in such approaches include the polyU polymerase PUP-2, the cytosine deaminase APOBEC, and adenosine deaminases ADAR and the engineered rABE. As the eventual readout employed in these approaches is deep sequencing, which has been extensively optimized for extraordinarily low inputs, we considered that such RNA tagging methods could be employed in early embryos via microinjection of mRNA encoding the RBP-editor fusion construct.

Here, we adopt REMORA ^1^, using the engineered RNA editor rABE, for use in the mammalian preimplantation embryo. We generated a series of 17 constructs fusing various RBPs to rABE (**Fig. 1A**), and characterized the targets of these RBPs at the four cell stage in mouse embryos to generate an Early Embryo RNA-RBP interaction (EERRi) dataset. We confirm expected behaviors, from subcellular localization to binding to known RNA motifs, for the majority of the RBPs under study, providing strong support for the validity of this approach. To identify target RNAs directly regulated by the RBP in question, we generated an independent RNA-seq dataset characterizing mRNA metabolism following siRNA-mediated knockdown of the RBPs under study, providing a robust dataset of regulatory functions of various RBPs in the mammalian preimplantation embryo. We focused follow up studies on Alyref, an RNA-binding protein involved in nuclear export, showing that Alyref plays a key role in target mRNA PolyA site usage, and that knockdown of *Alyref* mRNA causes embryo arrest prior to the morula stage. Overall, our results suggest a dynamic and developmentally vital RNA-RBP interactome, and provides a proof of concept for exploring a new arena of early embryonic studies.

**Figure 1.**
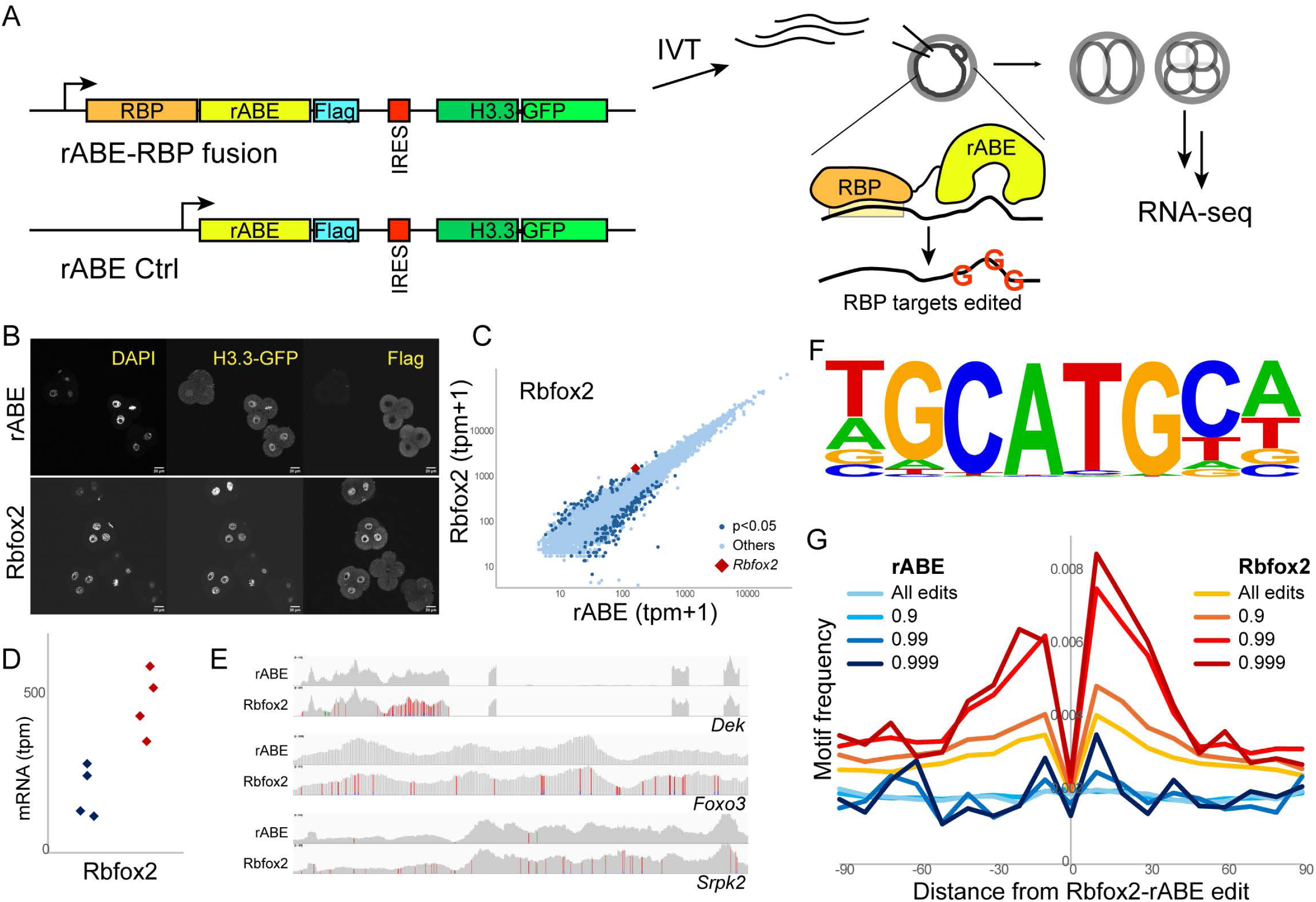
Proof of concept identification of RBP cargo in early mouse embryos. A) Schematic of REMORA approach in preimplantation embryos. In vitro transcription constructs were generated either with the engineered RNA Adenosine Base Editor (rABE) ^1^ alone, or with rABE fused in frame to an RNA Binding Protein (RBP) of interest. Both constructs carry an associated H3.3-GFP construct to allow visualization of successful zygote injections. mRNA produced by in vitro transcription is injected into control zygotes, driving the expression of rABE control protein or RBP-rABE fusions, resulting in A->I edits (which result in A->G “mutations” after reverse transcription during deep sequencing library preparation) in RNAs bound by the rABE-RBP fusion protein. B) Rbfox2-rABE exhibits expected nuclear localization. Confocal images show DAPI, H3.3-GFP, and Flag immunostaining, as indicated, for rABE only and rABE-Rbfox2-injected embryos. C) Modest effects of Rbfox2 overexpression on global mRNA abundance. Scatterplot shows median mRNA abundance for the four rABE-injected embryo pools (batch-corrected DESeq mRNA abundance + 1), y axis shows median abundance for Rbfox2-injected embryos. *Rbfox2* abundance is shown as a red diamond, with dark blue dots showing significantly-altered mRNAs (padj <= 0.05, log2FC >= 1). D) Dot plot of *Rbfox2* mRNA abundance for rABE-only embryos (left, blue) and rABE-Rbfox2 embryos (right, red). E) Examples of rABE-directed edits over high-confidence Rbfox2 targets. F) Sequence logo for sequences enriched surrounding high confidence rABE-Rbfox2 editing sites. G) Known Rbfox2 motifs are enriched surrounding rABE-Rbfox2 edit sites. Sequences were aligned over rABE only or rABE-Rbfox2 edits, called with the indicated stringency: all edits, P>0.9, 0.99, or 0.999. Y axis shows Rbfox2 motif frequency at varying distances (x axis) from edit sites.

## RESULTS

### Proof of concept mapping of RBP cargo preimplantation mouse embryos

We set out to characterize RNA binding protein (RBP) cargos in the early mouse embryo, adopting REMORA – <u>R</u>NA-<u>e</u>ncoded <u>mo</u>lecular <u>r</u>ecording in <u>a</u>denosines ^1^ – in which an engineered RNA adenosine base editor (rABE) is fused to the RBP of interest, leading to A->I edits (which are then captured as A->G “mutations” in downstream deep sequencing datasets) on RNAs associated with the RBP of interest. As a proof of concept, we sought to recapitulate the binding behavior for Rbfox2, which has a well-defined RNA binding motif and whose RNA cargos have been intensively characterized in multiple prior studies in a variety of cell types ^1,30,31^. In order to express REMORA constructs in preimplantation embryos, we generated an rABE-Rbfox2 fusion in a T3 in vitro transcription vector to allow mRNA production for zygote injections (with an IRES-H3.3-GFP cassette to visualize successful injections), along with matched rABE-only constructs to control for overall mRNA abundance and for any sequence specificity of the untargeted rABE enzyme (**Fig. 1A**). In addition, all proteins surveyed throughout this study were produced with a C-terminal Flag tag, enabling us to visualize the subcellular localization of the fusion protein (**Fig. 1B, Supplemental Fig. S1**).

We generated zygotes via in vitro fertilization (IVF), and two hours after removing sperm (six hours after beginning IVF) we injected ∼150 fL of 100 nM in vitro-transcribed mRNA into zygotes. We collected embryos 48 hours post-fertilization, selecting only those injected embryos that had reached the four cell stage. We then pooled ∼5 embryos per replicate for SMART-Seq3 ^32^ sequencing, targeting 3-6 replicates per condition; for this proof of concept, we generated 4 replicates each for the rABE control and Rbfox2 fusions. After mapping reads to the mouse transcriptome, edits were quantitated using the SAILOR pipeline ^33^, assigning each putative edit site a probability of being a true edit (vs. sequencing errors) based on the fraction of reads across that nucleotide exhibiting A->G mutations. Below, we compare rABE and Rbfox2 datasets using all editing calls, or edits selected for increasing levels of stringency (P>0.8, 0.9, 0.99, or 0.999).

Initial analyses confirmed that the Rbfox2-rABE fusion did not dramatically perturb the course of early development. Rbfox2-rABE-injected embryos attained the four cell stage at equivalent rates to rABE controls (81% vs. 84%). The Rbfox2 fusion protein localized to the nucleus (**Fig. 1B**), consistent with its expected localization, and our RNA-seq data revealed a modest ∼2-fold increase in *Rbfox2* RNA levels (**Figs. 1C-D**), indicating that the fusion RNA is present at comparable levels to the endogenous *Rbfox2* mRNA. This overexpression resulted in modest changes to gene expression in these embryos, with 203 genes exhibiting 2-fold up- or down-regulation at the four cell stage (**Fig. 1C, Supplemental Fig. S2, Supplemental Table S1**). Moreover, because rABE edits are calculated as a fraction of mRNA reads, any fusion protein effects on gene expression are internally controlled and should have little to no effect on the identification of bona fide Rbfox2 target transcripts.

**Fig. 1E** shows several examples of transcripts exhibiting extensive Rbfox2-specific editing, with a scattering of edits observed in the rABE-only control contrasting with scores of significant edits in the Rbfox2-rABE-injected embryos. Overall, we find a ∼3-10 fold increase in the number of rABE edits in the Rbfox-rABE dataset compared to the untargeted control: ∼41,200 vs ∼14,000 edits per dataset for all unselected edits, dropping to 2334 vs. 256 edits for the most stringent (P > 0.999) editing sites (**Supplemental Fig. S3A**). Rbfox2 edits were enriched in mRNA 3’ UTRs, with this 3’

UTR enrichment increasing with increasing stringency (**Supplemental Fig. S3B**), consistent with the known binding locations for Rbfox2 ^34^. Given the well-established binding motif for Rbfox2, we next searched for enriched sequence motifs surrounding the Rbfox2-rABE edit sites, recovering a good match for the known Rbfox2 binding motif (**Fig. 1F**). Rbfox2 motifs were strongly enriched within ∼40-50 nt of the Rbfox2-rABE edits, with increasing enrichment observed with more stringent edit calls (**Fig. 1G**). In contrast, we find no enrichment for Rbfox2 motifs surrounding the rABE-only edits, as expected. Similar to the pattern of 3’ UTR edit enrichment increasing with the stringency of edit calls, we find that the greatest improvement in motif enrichment is found moving from P>0.9 to P>0.99 stringency for the editing calls; we therefore use the P>0.99 edit datasets for most analyses below unless otherwise specified.

Taken together, these data confirm that REMORA can be used to characterize RBP-associated RNA cargos in mammalian preimplantation embryos, motivating us to expand our studies to survey a number of additional RBPs.

### A survey of 17 RBPs in the early embryo

Confident that REMORA can provide specific information on RBP binding targets in early embryos, we generated an additional 16 RBP-rABE fusion constructs for analysis. Several RBPs were chosen as additional positive controls to provide further validation for our approach and pipeline, along with a number of RBPs with known roles in the early embryo, as well as a handful of underexplored RBPs chosen based on their expression at this developmental stage. We confirmed expected localization patterns for all 17 RBP fusion proteins (**Supplemental Fig. S1**), and documented effects of the

RBP-rABE mRNA on mRNA abundance (**Supplemental Fig. S2, Supplemental Table S1**). In general, we find between 100 and 500 significantly-misregulated genes for most RBP-rABE fusions, with two major exceptions. Specifically, both Elavl1-rABE and Ybx3-rABE-injected embryos exhibited substantial alterations in development, with only ∼36% of Elavl1-injected embryos reaching the four cell stage, and all (40/40) Ybx3-injected embryos arresting in the two-cell stage. In the case of Elavl1, enough four cell embryos could be collected for further analysis. In the case of Ybx3, given the two cell stage arrest, we collected a 30 hour post-fertilization dataset to define Ybx3 targets. In both of these scenarios, we identified much more dramatic gene regulatory changes than observed for the other RBPs in this study (**Supplemental Fig. S2**), consistent with the developmental changes driven by these constructs. We therefore urge some caution regarding the interpretation of the Elavl1 and Ybx3 datasets, although as noted above the internally-controlled nature of the REMORA approach should largely control for the altered physiological state of the injected embryos.

Overall, our set of RBPs exhibited a ∼100-fold range in the number of edited targets, with between ∼100 edits (P>0.99) per dataset for Mrps22 ranging up to ∼30,000 edits per dataset for Ybx3 (**Fig. 2A, Supplemental Table S2**). Altogether, roughly half of the RBPs investigated exhibited increased RNA editing activity compared to the untagged rABE control. Importantly, the scarcity of RNA targets was expected for many of the RBPs with few identified targets. Mrps22 is a component of the mitochondrial ribosome; given the minimal number of transcripts expressed from the mitochondrial genome, and the rare use of polyA tailing on these transcripts, we did not expect to observe many targets for Mrps22. Nonetheless, despite the inefficient capture of mitochondrial RNAs by the PolyA-dependent SMART-Seq protocol, our dataset included these transcripts at low abundance, and Mrps22 was the only RBP to drive editing of mitochondrial transcripts (**Fig. 2B, Supplemental Fig. S3C**), providing further confidence in our approach. Similarly, Slbp is expected to bind only to histone-encoding mRNAs ^35^, which carry a unique 3’ UTR and are not typically polyadenylated, and thus are not expected to be present in these datasets. Indeed, we did not capture the canonical histone mRNAs – tpm values were 0 across the dataset for the majority of canonical histone mRNAs – and were thus unable to test for the expected Slbp-driven editing of histone mRNAs. Cnot8 is a member of the CCR/NOT RNA deadenylation complex; here, the modest editing observed is presumably a result of Cnot8-bound targets being deadenylated and degraded and thus unavailable to be captured for analysis. Finally, Carm1/Prmt4 is a histone arginine methylase and does not carry a known RNA binding domain, but was chosen for this study based on reports that *Neat1* RNA plays a role in Carm1 localization ^20^; however, we find no substantial Carm1-dependent edits in *Neat1* (not shown), perhaps reflecting the fact that Carm1 is thought to be recruited to *Neat1* via adaptor proteins and thus is not localized in immediate molecular proximity to this RNA. Somewhat more surprising was the low editing efficiency for Rps2 and Rps3, especially given the reported success of Rps2 and Rps3 fusions to APOBEC in the “Ribo-STAMP” protocol ^29^. We speculate that the failure of Rps-rABE fusions here may reflect poor positioning of the N-terminal rABE, which seems likely to affect tagged Rps2/3 association with the 80S ribosome; perhaps optimizing linker sequence or length, or altering the tagged side of the ribosomal protein, would improve the success of this approach.

**Figure 2.**
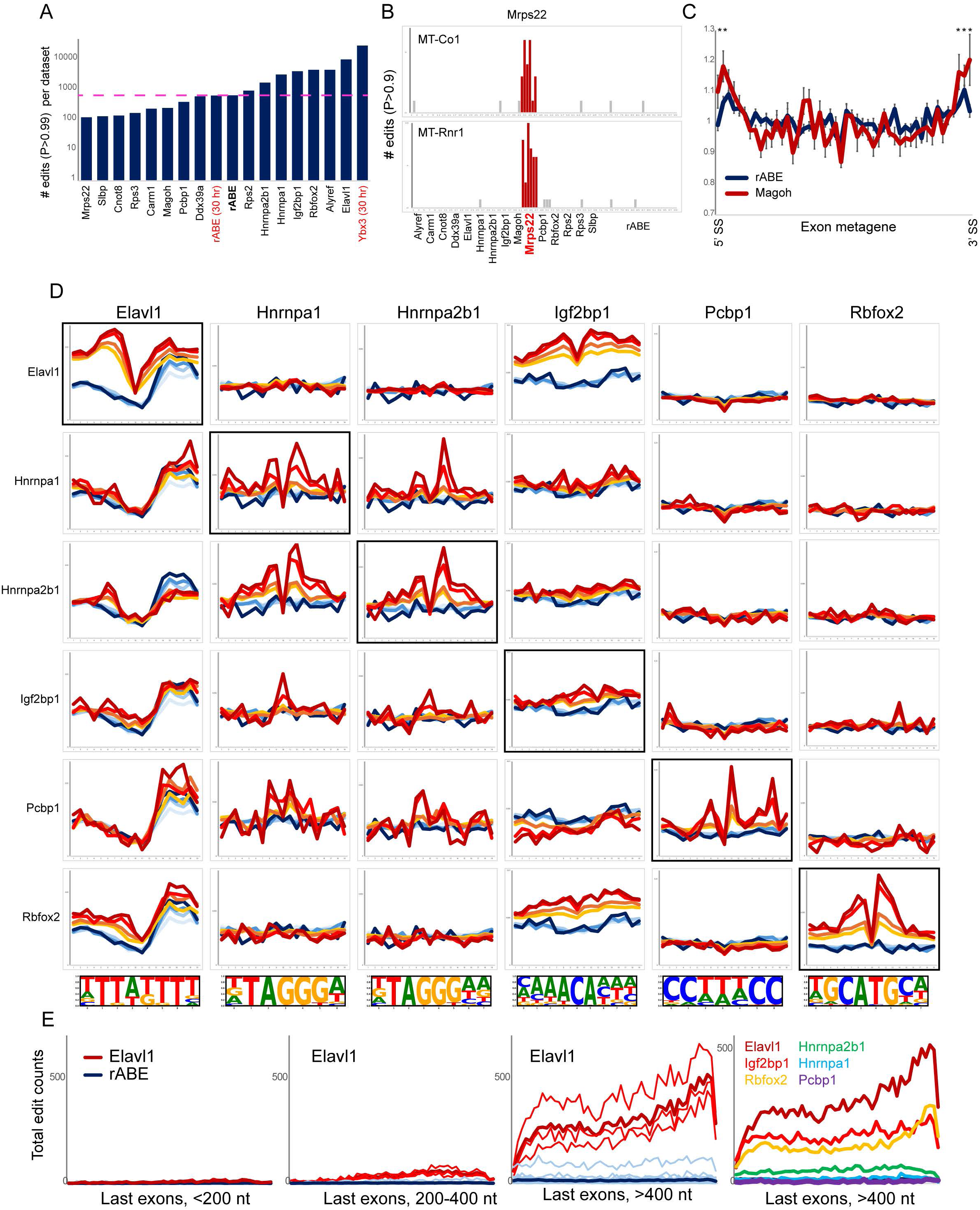
Overview of the 17 RBP dataset. A) Average number of edits (P>0.99) for the indicated RBPs, or untagged rABE controls. All datasets were collected at 48 hours post-IVF with the exception of the matched rABE and Ybx3 datasets collected at 30 hours, indicated in red. Dotted line shows average edit numbers for untargeted rABE. B) Number of edits (p>0.9) for two mitochondria-encoded mRNAs, showing editing exclusively in the Mrps22 dataset (indicated in red). C) Metagene shows edit frequency across length-normalized exons, showing significant (**, ***: p <0.01, 0.001 respectively) editing at exon-exon junctions in the Magoh dataset. D) Motif frequency for known motifs for the indicated RBPs (motifs shown at the bottom of each column) as a function of distance from RBP-rABE edits. Data are visualized as in Fig. 1G. E) Association of specific RBPs with long 3’ terminal exons. Left three panels show Elavl1 binding to short, medium, and long final exons, as indicated (thin lines show individual replicates, thick line shows average). Rightmost panel shows average editing activity for six relevant RBPs.

To validate our approach, we next turn to a number of RBPs which were chosen as positive controls for this study based on clear predictions for their RNA cargos. **First**, as noted above, despite driving the fewest edits of any RBP in this study, Mrps22 was, as expected, the only RBP to direct substantial editing of mitochondrial RNAs. **Second**, Magoh is a component of the exon junction complex; consistent with this function, we find edits enriched at exon-exon junctions in spliced mRNAs, and depleted from introns and 5’ and 3’ untranslated regions (**Fig. 2C**). **Third**, although the modest number of edits observed for Rps2 did not suggest successful incorporation into the ribosome, at the very least the cytoplasmic localization of the fusion would be expected to confine Rps2-rABE edits to spliced mRNAs; as expected, Rps2 editing datasets were enriched for exons and depleted of introns and untranslated noncoding RNAs (**Supplemental Fig. S3D**). Among our positive controls, only Slbp – which is expected to bind to the histone stem loop element in the 3’ UTR of histone mRNAs – failed to exhibit the expected behavior, as canonical histone-encoding mRNAs were not captured in our SMART-Seq3 libraries.

**Finally**, we characterized editing patterns for RBPs with well-characterized sequence motifs, as shown above for Rbfox2 (**Fig. 1G**). Overall we find good enrichment of motifs for Elavl1, Hnrnpa1, Hnrnpa2b1, and Pcbp1 surrounding the editing sites identified here (**Fig. 2D**). Moreover, there was little to no enrichment of inappropriate RBP motifs in any given dataset, with two exceptions. First, because of the near-identity of their target motif sequences, we observe enrichment for both Hnrnpa1 and Hnrnpa2b1 motifs in both proteins’ datasets. Second, although we observe modest but significant enrichment for the reported motif for Igf2bp1 surrounding

Igf2bp1 edits, more dramatic enrichment is seen for this motif surrounding Elavl1 and Rbfox2 edits (**Fig. 2D**). The strong enrichment for the Igf2bp1 motif presumably reflects the role for Igf2bp1 in recruiting Elavl1 to target transcripts ^36^; while a similar role in recruiting Rbfox2 has not, to our knowledge, been previously described, the modest enrichment of Rbfox2 edits around this motif may instead reflect the shared functions of these proteins as readers of the covalent RNA modification 6-methyl adenosine (m6A). Indeed, a number of the RBPs under study here have previously been implicated as m6A readers. To further explore this connection, we compared RBP cargos to a prior dataset characterizing m6A levels in early embryos ^37^ (**Supplemental Fig. S3E**). In addition, as m6A is typically enriched on the final exons of mRNAs, particularly on longer exons ^38,39^, we also plotted edits for six RBPs over final exons binned according to length (**Fig. 2E**). Consistent with their known preference for m6A-modified RNAs, we find strong enrichment for Elavl1 ^40^, Igf2bp1 ^41^, and Rbfox2 ^42^ over long exons (**Fig. 2E**) and at previously-characterized m6A-marked transcripts in early embryos (**Supplemental Fig. S3E**), as well as moderate but significant enrichment for Hnrnpa2b1 (which binds to m6A – ^43^) but no enrichment in either analysis for Hnrnpa1 (which does not) or Pcbp1.

Taken together, these analyses further support the ability of REMORA to identify RNA targets for a subset of RNA binding proteins in early mammalian embryos.

### Effects of RBP binding on target RNA metabolism

To determine the functional consequences of RBP binding on the life cycle of target RNAs, we carried out siRNA-mediated knockdowns of each of the RBPs in our primary dataset and characterized effects in resulting four cell stage embryos on mRNA abundance and structure (splicing, alternative TSS or polyadenylation sites, etc.) by SMART-seq (**Supplemental Table S3**). We achieved good knockdown efficiency, with the majority of targeted RBPs exhibiting at least 10-fold lower abundance under our KD conditions (**Fig. 3**). Given the role for RBPs in all aspects of post-transcriptional RNA metabolism, from splicing to RNA stability, we analyzed our data not only for mRNA abundance but also for altered patterns of splicing and polyadenylation ^44,45^. Under these conditions, we find a wide range of effects on mRNA abundance, splicing, and 3’ end processing across our set of RBPs; for instance, focusing on mRNA abundance, we document a range from 35 significant differentially-expressed genes (DEGs) for *Rbfox2* KD, up through 1189 DEGs for *Alyref* KD (median of 398 DEGs across the dataset), with relatively few differentially-expressed genes following knockdowns of *Rbfox2* (35 DEGs), *Elavl1* (66), and *Hnrnpa1* (60). These RBPs presumably represent either proteins that are redundant with other proteins not perturbed here, proteins with functions in RNA biology that are confined to specific conditions (stressors etc.) or not measured here (nucleotide modifications, etc.), or – in our view most likely – relatively stable proteins which would be minimally affected at the protein abundance level despite the robust RNA knockdown efficiencies here (**Fig. 3**). In other words, as in any knockdown study, it is important to be aware that differing protein stabilities will mean that the robust RNA knockdown efficiencies will nonetheless result in variable effectiveness in removing RBP function.

**Figure 3.**
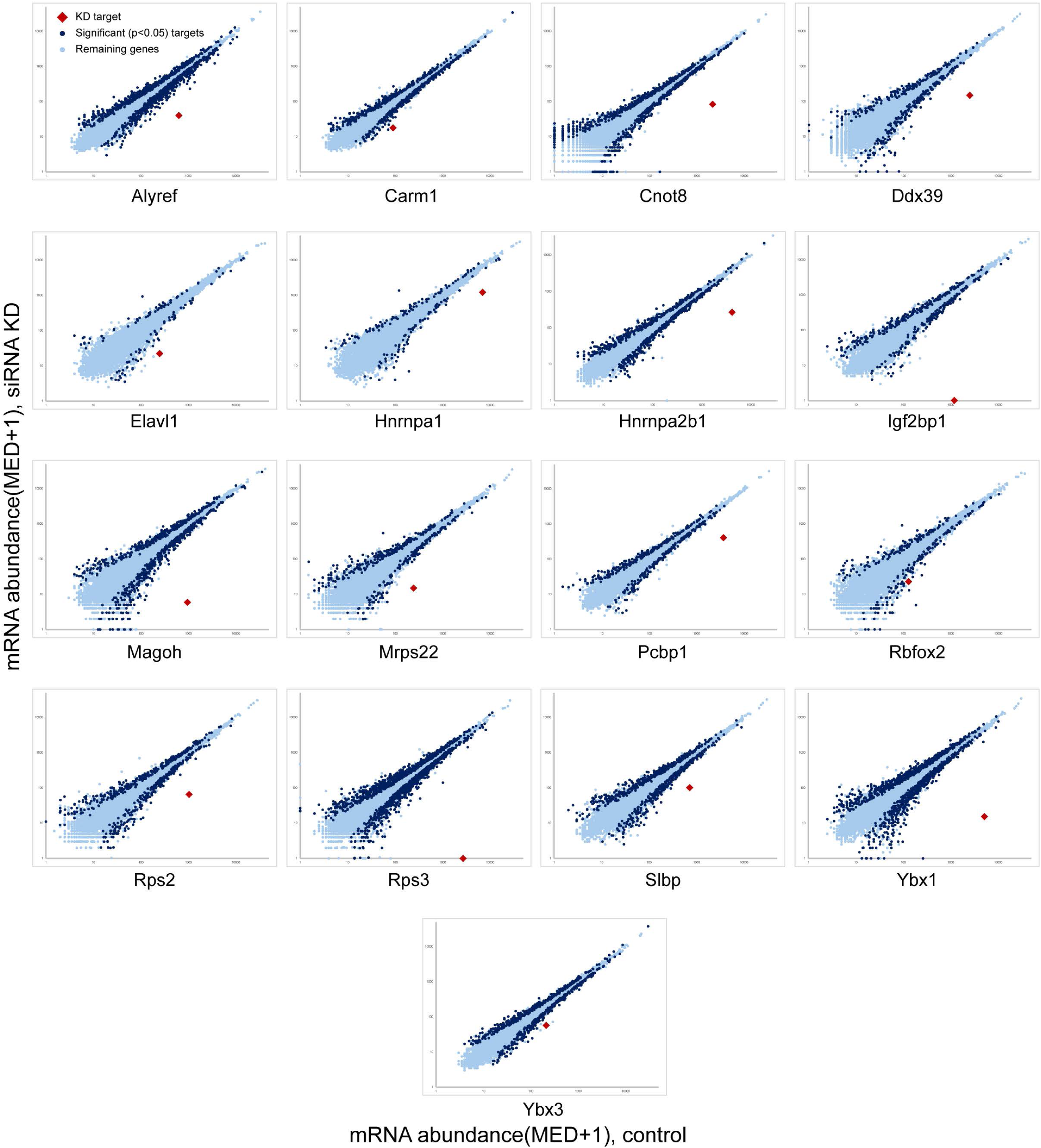
RNA-seq data following RBP knockdowns. Scatterplots show changes in mRNA abundance following RBP knockdown (y axis), relative to matched control knockdown data (x axis). The targeted mRNA, indicated in red, is typically downregulated ∼5-10-fold.

In order to focus analysis on functional interactions between RBPs and bound RNAs, we searched for significant overlap between genes misregulated following RBP knockdown, and transcripts directly bound by a given RBP in our EERRi dataset. **Fig. 4** shows the five RBPs with the most significant overlap (p < 10^−3^) between RNA cargoes and RNAs affected at the level of mRNA abundance. Similar analyses are shown for alternative splicing in **Supplemental Fig. S4** and **Fig. 5**, and for alternative polyadenylation site usage in **Fig. 6** and **Supplemental Fig. S5**. These analyses identify a cohort of transcripts likely to represent direct targets of RBP-mediated regulation in the preimplantation embryo, highlighting the diversity of roles for RBPs in gene regulation in the mammalian embryo.

**Figure 4.**
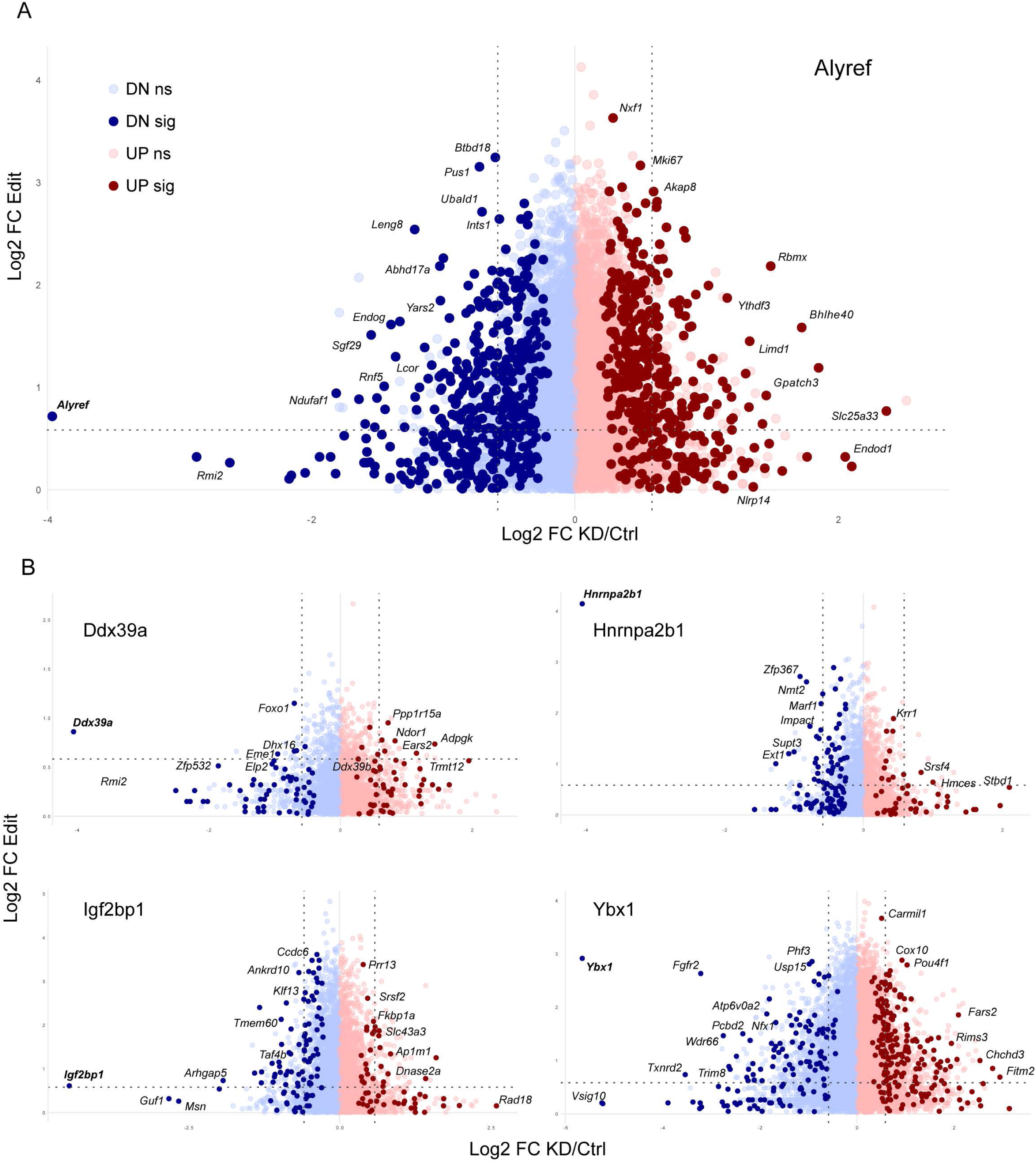
Significant effects of RBP knockdowns on RBP cargo RNAs. A) Scatterplot shows RNAs bound by Alyref on the y axis, calculated as the log2 fold change in Alyref-rABE edits divided by rABE-only edits (both values +1 to reduce noise and avoid dividing by zero); dataset only shows RNA cargos with more Alyref-rABE edits than rABE control edits. X axis shows changes in mRNA abundance following Alyref knockdown, with significant (padj < 0.05, DESeq2) differentially-expressed genes shown in darker blue/red. B) Overlap between RBP binding (y axis) and gene expression changes following RBP knockdown (x axis) for four additional RBPs with significant overlap between RNAs bound by the RBP and RNAs significantly affected by RBP knockdown.

**Figure 5.**
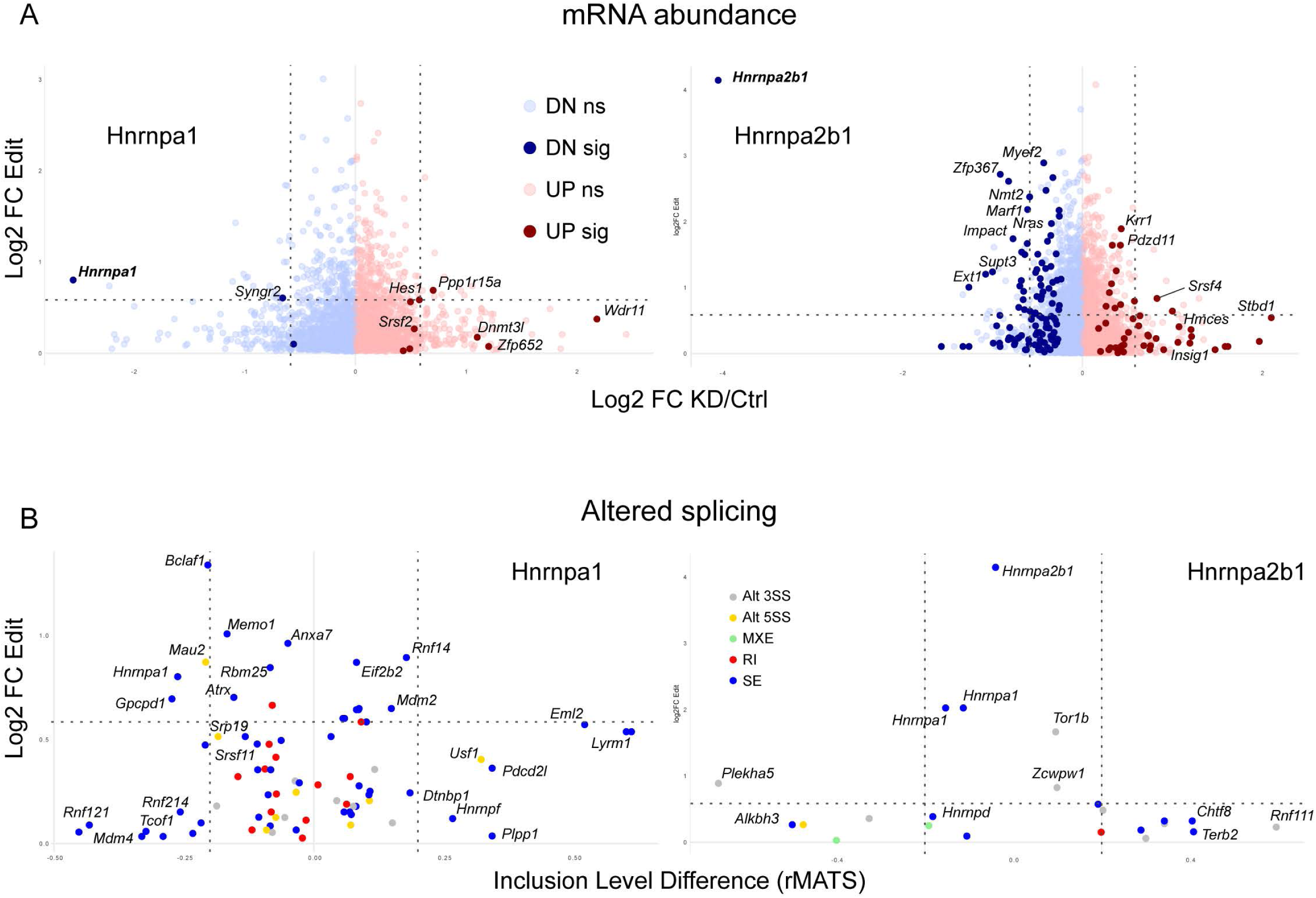
Distinct functions for Hnrnpa1 and Hnrnpa2b1 in RNA metabolism. Distinct functions for Hnrnpa1 and Hnrnpa2b1 in embryonic gene regulation. (A) shows changes in mRNA abundance following RBP knockdown, as in Fig. 4. (B) shows similar data for splicing changes (x axis) as called by rMATs ^44^. Alt3SS and Alt5SS: altered 3’ or 5’ splice sites; RI: retained intron; SE: skipped exon; MXE: mixed effects. X axis shows extent of difference in splicing inclusion: alternate exon (or 5’ splice site usage, etc.) level inclusion in KD - Ctrl.

**Figure 6.**
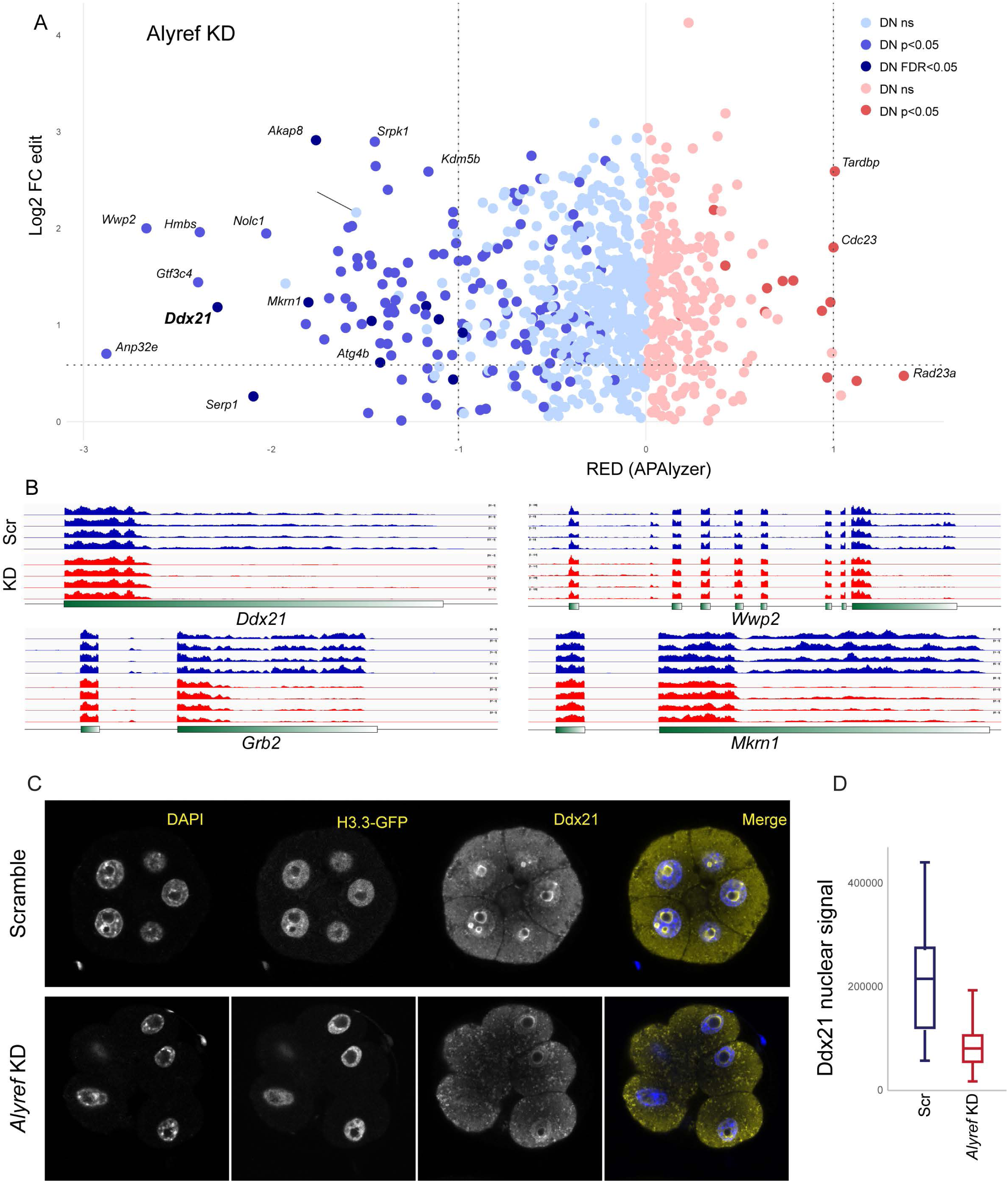
Alyref controls stability of specific mRNA isoforms. A) Changes in 3’UTR length following *Alyref* KD. X axis shows relative expression difference (RED, output from APAlyzer ^45^) between longer and shorter 3’ UTR isoforms: positive values indicate longer 3’ UTRs, negative values indicate shorter UTRs. Y axis shows Alyref-rABE edits, as in **Figs. 4-5**. Dots are colored by significance of change in 3’ UTR length: p>0.05 (ns), p<0.05 but FDR > 0.05, and FDR < 0.05. B) Examples of mRNAs exhibiting shorter 3’ UTRs following *Alyref* KD. 4 representative rABE only controls (out of 26) are shown for simplicity. C) Images of control (injected with scrambled siRNAs) or *Alyref* knockdown embryos at 54 hrs post-fertilization (eight cell stage). Images show DAPI, H3.3-GFP, and Ddx21, as indicated. Note the loss of Ddx21 nuclear fluorescence following *Alyref* KD. D) Quantitation of Ddx21 nuclear staining in control (n=55) and *Alyref* KD (n=77) embryos. Boxplots show maximum, 75^th^ percentile, median, 25^th^ percentile, and minimum.

Our data highlight the distinct functions of Hnrnpa1 and Hnrnpa2b1 ^46–48^ in gene regulation in the embryo. These proteins bind to similar RNA sequence motifs (**Fig. 2D**), and both proteins have been implicated in a range of functions in RNA metabolism, most notably including mRNA splicing ^49,50^ but extending to mRNA stability ^51^, localization ^52^, and even small RNA loading into exosomes ^53^. Here, we find a notable divergence in functions of Hnrnpa1 and Hnrnpa2b1 in mammalian embryos, as we find that Hnrnpa2b1, but not Hnrnpa1, significantly affects mRNA abundance (**Fig. 5A**). In contrast, while both Hnrnpa1 and a2b1 exert significant effects on mRNA splicing, Hnrnpa1 drives far more significant and extensive effects on mRNA spicing patterns (**Fig. 5B**). These analyses highlight divergent functional consequences of two related RBPs with similar mRNA binding behavior.

We finally turn to effects of RBPs on mRNA 3’ UTR length. Although five RBPs exhibited significant overlap between edited RNA cargos and transcripts exhibiting a 3’ UTR length change following RBP knockdown at a nominal (uncorrected) p value of 0.05 (**Supplemental Fig. S5**), only *Alyref* KD drove significant effects on 3’ UTR lengths at the more stringent FDR < 0.05 for UTR length changes (**Fig. 6A**). Notably, *Alyref* knockdown systematically drove shortening of mRNA 3’ UTRs: for affected transcripts, mRNA coverage data show that control embryos (as well as unrelated KDs, not shown) appear to express a predominant short mRNA isoform along with a subpopulation of mRNAs with longer 3’ UTRs, and this longer mRNA isoform is specifically lost following *Alyref* KD (**Fig. 6B**).

Among the transcripts exhibiting Alyref-driven changes to 3’ UTR length, we noted that Ddx21 is essential for embryo development to the blastocyst stage ^54^. To explore the functional consequences of the observed shortening of the *Ddx21* 3’ UTR, we quantitated Ddx21 protein levels in control and knockdown embryos, finding significantly decreased levels of Ddx21 protein following *Alyref* KD (**Figs. 6C-D**).

Consistent with the loss of Ddx21 production, we confirm that *Alyref* KD embryos exhibit delayed compaction at the 8-cell stage (**Supplemental Figs. 6A-B**), and – consistent with previous reports ^24^ – do not progress normally to the blastocyst stage, with weak or absent cavitation at 96 hours post-fertilization (**Supplemental Fig. S6C**).

## DISCUSSION

The preimplantation embryo represents a unique system for gene regulation, with dramatic upheaval affecting everything from genome packaging to post-transcriptional gene regulation. Although this developmental window has been challenging to study in mammals thanks to the limited availability of samples, advances in genomic technologies have driven remarkable progress in recent years in our understanding of chromatin packaging and transcription. Here, we extend these efforts to provide a proof of concept for exploring post-transcriptional gene regulation by RNA binding proteins.

We adopted the REMORA approach to fusing RNA editors to an RNA-binding protein of interest, expressing 17 RBP-rABE fusions (along with the untargeted rABE control) in preimplantation embryos to label the RNA targets of RBPs of interest.

Benchmarking our data against known features of these RBPs, we estimate that we accurately assay the binding for roughly half of the RBPs under study. Our data provide an atlas of target RNAs in the preimplantation embryo for Alyref, Ddx39a, Elavl1, Hnrnpa1, Hnrnpa2b1, Igf2bp1, Mrps22a, Pcbp1, Rbfox2, Ybx1, and Ybx3, while our data for Mrps22 and Magoh are also consistent with expectations albeit with poor coverage. In contrast, we have low confidence in our data for Rps2, Rps3, Cnot8, Slbp, and Carm1; the failure of the RPGs may reflect accessibility issues with the N-termini of these proteins, while the latter three presumably failed as Slbp is not expected to target polyadenylated transcripts, Carm1 may have no direct RNA targets, and Cnot8-bound targets may be destroyed before being available for sequencing. Altogether, along with a recent study from the Conine lab adopting REMORA to characterize targets of Ago2 ^55^, our data open the door to systematic analysis of RNA binding proteins in early development.

We also generated a significant resource of gene expression changes caused by knockdowns of the 17 RBPs under study here. These data are affected by typical caveats of knockdown studies; although our knockdowns very efficiently depleted mRNAs (**Fig. 3**), any stable pre-existing proteins would continue to function in RNA metabolism and blunt the impact of the knockdowns. Nonetheless, we observed strong effects for over half of our knockdowns, and for roughly half of the knockdowns under study we identified significant overlap between RNA targets identified by REMORA and RNAs exhibiting changes in abundance, splicing, or polyadenylation (**Figs. 4-6**), identifying strong candidates for direct RBP-mediated gene regulation in the embryo.

Among the roles for RBPs in embryonic gene regulation, we highlight pleiotropic roles for the nuclear export adaptor Alyref in mRNA metabolism in embryos, with effects on mRNA abundance, splicing, and, most dramatically, on control of 3’ UTR length (**Fig. 6**). Best-known as a member of the transcription export (TREX) complex ^56^, Alyref also plays diverse roles in mRNA metabolism from control of splicing to maintenance of genome stability via prevention of R loop formation ^57^. Alyref is loaded onto mRNAs co-transcriptionally via the exon junction complex ^58^, and multivalent binding between Alyref and EJCs has been proposed to help compact ribonucleoprotein complexes ^59^. In addition to its loading over coding sequences mediated by the EJC, Alyref is also recruited onto transcripts via interactions with the 3’ end processing machinery ^60,61^, and in yeast was shown to modulate 3’ mRNA end processing ^60^, although Yra1 (yeast Alyref) depletion lead to both shortening and lengthening of 3’ UTRs at different transcripts, contrasting with the preferential loss of long mRNA isoforms documented here. Overall, we confirm pleiotropic roles for Alyref in the preimplantation embryo, with *Alyref* KD driving changes to mRNA abundance, splicing, and 3’ UTR usage. It will be interesting in future studies to define how Alyref binding in different contexts drives distinct functional outcomes; we note here that different binding locations for Alyref correlate with distinct outcomes, as Alyref binding over coding sequences was correlated with transcripts upregulated following *Alyref* KD, whereas Alyref binding to 3’UTR and CDS was associated with downregulation in *Alyref* knockdowns (not shown). Finally, we define one functional target of Alyref – Ddx21 – required for normal development, providing one possible explanation for the developmental defects seen in *Alyref* KD embryos.

## ACKNOWLEDGEMENTS

We thank S. Floor for generous gift of the rABE construct, and G. Yeo, E. Kofman, and B. Yee for assistance with the SAILOR computational pipeline. Work was funded by the Lalor Foundation (PB) and NIHR01HD099816 (PB, OR).

## MATERIALS AND METHODS

### Ethics statement

Animal husbandry and experimentation were reviewed, approved, and monitored under the University of Massachusetts Chan Medical School Institutional Animal Care and Use Committee (IACUC protocol no. 201200029).

### Animal husbandry

Female and male FVB/NJ mice at ages of about 8 and 10 weeks, respectively, were used in this study. The animals were maintained under controlled temperature and humidity conditions on a 12-hour light/dark cycle and fed control diets (AIN-93G pellets).

### Embryo production and culture

Female mice were superovulated by intraperitoneal injections with 7.5 IU of PMSG or 100ul of CARD HyperOva (Cosmo Bio USA KYD-010-EX-X5) followed by 7.5 IU of hCG ∼46 hours later. 12-14 hours thereafter, male and the superovulated females were sacrificed and dissected to isolate sperm and cumulus-oocyte-complex (COC), respectively. Sperm from the cauda epididymis and vas deferens was squeezed out into a 100ul drop of HTF (Tribioscience TBS8072) supplemented with 0.75mM methyl-β-cyclodextrin (Millipore-Sigma C4555) and covered with mineral oil (Millipore-Sigma M8410-500ML) in a 35mm culture dish, which was incubated at 37°C, 5% CO2 and 5% O2 for 30 minutes prior. After collection of sperm into the HTF drop, the dish was further incubated in the aforementioned condition for another hour to capacitate the sperm.

Simultaneously, COC was dissected out to preincubated drops of KSOM (Millipore-Sigma MR-106-D) and then moved to 50ul drops of preincubated HTF, usually one COC into one drop. Once capacitated, 5-8 μl of the sperm from the periphery of the HTF capacitation drop, was added to each 50ul HTF drop containing the COCs. This time-point was marked as the start of in vitro fertilization (IVF), and the plates were placed back in the incubator for 1.5 to 2 hours. Thereafter, the oocytes were moved to preincubated KSOM drops, washed by moving through few drops of the medium and then placed back into the incubator for 2 to 3 more hours for post-IVF recovery.

Fertilized oocytes were identified by appearance of two pronuclei i.e. zygotes and used for microinjections. Microinjections were carried out in M2 (Millipore-Sigma M7167) drops covered with mineral oil and under an inverted microscope (Zeiss Axio Observer) assembled with microinjection setup (Eppendorf; TransferMan 4r, CellTram Air 5176, FemtoJet 4i), a glass thermoplate (Tokai-Hit) at 37°C and occasionally aided by a MICRO-ePORE system (WPI). Microinjections were carried out using holding pipettes (Sunlight Medical SHP-100-35) and needles pulled from borosilicate glass pipettes with capillaries (Sutter Instrument BF100-78-10) using a Sutter Instruments P-97 micropipette puller. The injection mix consisting of the experimental RNA construct, usually coupled with a marker mRNA in the form of histone-GFP, is prepared in ultra-pure water (Thermo Fisher Scientific AM9932) and loaded into the needle using a microloader pipette-tip (Eppendorf 930001007). In batches of about 10 zygotes, the embryos were microinjected and then moved back to KSOM drops to be cultured. 28-30 hours after fertilization, that is at the late-2-cell stage, the embryos were checked for successful microinjection by visualization of a fluorescent nucleus and then cultured to the desired developmental stage.

### Molecular biology

The rABE and Rbfox2-rABE constructs used in this study were reported in Lin *et al.* ^1^ and the relevant plasmids were obtained from Addgene (#191383 and #191384). rABE and Rbfox2-rABE sequences, from these plasmids, were amplified out and cloned into a vector for in vitro transcription (IVT). Other candidate genes for this study were amplified from mouse ES cell and blastocyst cDNA and cloned so as to replace the Rbfox2 sequence in the IVT plasmid. The IVT plasmids are therefore of the following general design: T3 promoter-RBP-rABE-FLAG-IRES-histone-GFP. The constructs were designed and generated using NEB HiFi DNA Assembly kit (NEB E2621) and NEBuilder. The plasmids thus generated, were linearized using a restriction site downstream of the histone-GFP sequence, and mRNA was synthesized using mMESSAGE mMACHINE™ T3 Transcription Kit (Thermo Fisher Scientific AM1348), followed by poly(A) tailing (Thermo Fisher Scientific AM1350). The synthesized mRNA was then quantified, checked in an agarose gel and used at 100 nM concentration for microinjections. In case of siRNA microinjections, a histone-GFP mRNA was added in the injection mix as an injection marker. This mRNA too was synthesized as described above, but from a plasmid with only the histone-GFP sequence.

### Microscopy

The processing of embryos for confocal immunofluorescence microscopy was as per published protocol ^54^. The embryos were fixed in the dark with 4% paraformaldehyde (Santa Cruz Biotechnology sc-281692) for 20 minutes at room temperature.

Permeabilization was carried out by transferring the embryos to a 0.5% solution of Triton X-100 (Millipore-Sigma T8787) in phosphate-buffered saline (PBS; 10x PBS Fisher BP399-500) for 20 minutes at room temperature. All washes after the fixation, permeabilization, and antibody staining procedures were in PBS with 0.05% TWEEN 20 (Millipore-Sigma P9416) (PBST) by transferring the embryos through three wells of PBST or wells in 96-well flat-bottom plates, for a total of 20-30 minutes min at room temperature. Blocking and antibody staining was in 3% bovine serum albumin (BSA; Millipore-Sigma A7906) in PBST. For blocking, embryos were incubated for 30 min at 4°C before both primary and secondary antibody staining. For primary antibody staining (in blocking buffer), embryos were incubated overnight (∼16 h) at 4°C. Secondary antibody staining was carried out in the dark at room temperature for 70 min. The stained embryos were mounted with VECTASHIELD mounting medium containing DAPI (Vector Laboratories H-1200), placed on glass-bottomed 35 mm dishes (Avantor/Matsunami D35-14-1-U), and incubated at 4°C for at least 30 min in the dark before imaging. Confocal images were acquired using a Nikon confocal system (A1RHD25).

### Image analysis

Quantification and analysis of the fluorescence images was as per published protocol ^54^ and primarily using Fiji (ImageJ) ^62^. To quantify the embryo as a whole, images from the channel pertaining to the required wavelength were analyzed as follows: Image > Stacks > Z Project. The projection-type “Sum Slices” was used to obtain a Z-projected image for the entire embryo. The levels of nuclear proteins were quantified as the fluorescence intensity per cell nuclei, and the cells were defined as inner or outer cells based on location of the nuclear DAPI immunofluorescence. The measurements for all of the above were set as follows: Analyze > Set Measurements, and the following options were chosen “Area”, “Mean gray value”, and “Integrated density”. Using the “Polygon selections” tool, an area encompassing the Z-projected embryo image (for whole embryo) or individual cell nuclei (for nuclear proteins) was demarcated, and the aforementioned measurements were recorded. The selected area was then moved to encompass an area excluding the embryo or cell nucleus, and background measurements were recorded. This process was continued for all the embryos analyzed under both control and experimental conditions, and the results were transferred to a spreadsheet for further calculations. The corrected fluorescence (cf), in arbitrary units, was measured for each embryo as follows: cf = integrated density – (area of selected cell × mean fluorescence of background readings). The calculated cfs are plotted in scatterplots, with the means and standard deviations marked.

### Library (from embryos)

Library preparation from embryos was using Smart-seq3 protocol ^32^, with necessary modifications. In brief, each pool of ∼5 embryos at desired developmental stage were collected in 5 μl of 1x TCL buffer (Qiagen 1070498) supplemented with β-mercaptoethanol (Millipore-Sigma M6250) at 1% concentration, in 8-strip PCR vials, which could then be stored at −80°C until next step. Immediately prior to mRNA isolation step of library preparation, another 5 μl of the TCL buffer was added to each sample and then incubated at room temperature for 10 minutes. 2.2 volumes of RNAClean XP beads (Beckman Coulter A63987), completely resuspended and at room temperature, was added to each sample and mixed thoroughly. This was then incubated at room temperature for 10 minutes, followed by a quick spin and then placed on a magnetic rack for another 5 minutes. Without disturbing the beads, the clear supernatant was carefully pipetted out, the beads were then washed thrice with 75 μl of freshly prepared 80% ethanol. After the last wash, the vials were left open to dry, but only until the beads appeared glossy. Excess drying of the beads results in poor library quality. The vials were then removed from the magnetic rack and 6 μl of Smart-seq3 lysis buffer was added to each, and the beads thoroughly resuspended in it. The vials were then placed in a thermocycler to carry out the cell lysis reaction, for 10 minutes at 72°C. The vials were placed on ice and 2 μl of reverse transcription mix was added to each and mixed thoroughly. This was followed by reverse transcription reaction in a thermocycler. Whole transcriptome amplification of the cDNA was carried out by adding 12 μl of the whole transcription amplification mix to it and running the 20-cycle amplification reaction in a thermocycler. The amplified library was then cleaned with either 0.65 volume of SPRIselect beads (Beckman Coulter B23318) or 0.9 volume of AMPure XP beads (Beckman Coulter A63881). Required volume of beads at room temperature and completely resuspended, was added to the reaction mix from amplification step, and mixed thoroughly. The vials were then incubated at room temperature for 5 minutes, followed by a quick spin and another 5 minutes on a magnetic rack. The supernatant was then pipetted out without disturbing the beads, and the beads were washed thrice with 100 μl of freshly prepared 80% ethanol. As previously, the beads were dried and then resuspended in 20 μl of TE buffer. 1 μl of this stock library was used for quantification using 1x dsDNA HS assay (Thermo Fisher Scientific Q33231) and a few samples were also used for fragment analysis as a quality-control of the libraries prepared. Following which, 1ul of the stock library was diluted to a working concentration of 0.5 ng/μl with TE buffer and used for the next steps. 1ul of the 0.5 ng/μl library was used for tagmentation reaction using tagmentation reagent from the Illumina Nextera XT DNA library preparation kit (FC-131-1096) at 55°C for 10 minutes, cooled to 4°C and then immediately followed by mixing with transposase stripping buffer (NT buffer or 0.2% SDS) and incubation at room temperature for 5 minutes. This was then followed by indexing with Nextera XT v2 indices (FC-131-2001/2/3/4) and fragment amplification. The final, amplified, individual libraries were combined in a 1.5ml low-bind tube and to it 0.8 volume of SPRIselect beads was added, mixed well and incubated at room temperature for 5 minutes. This was followed by a quick spin and 5 more minutes of incubation on a magnetic rack. As in the prior bead purification steps, the clear supernatant was removed without disturbing the beads, and the beads were then washed twice with 500 μl of 80% ethanol, dried and resuspended in 25-30 μl of TE buffer. This was followed by another 2 minutes incubation at room temperature and then 5 minutes on a magnetic rack. The supernatant, final library, was pipetted out and 1ul of this was used to quantify using 1x dsDNA HS assay and also subjected to fragment analysis to check its quality and measure average fragment size of the library. The pooled library was first diluted to 4 nM with TE buffer and then processed via denaturation and further diluted to 1.8-2 pM using HT1 hybridization buffer as is required for the Illumina NextSeq 500/550 system and sequenced using high-output v2.5 kits (Illumina 20024906). The Ybx1-rABE SMART-seq3 libraries were sequenced at Admera Health with NovaSeq X Plus 2×150 platform.

### Data analysis

RNA-sequencing libraries were demultiplexed with UMI-tools, strand-wise aligned with STAR ^63^ and mapped to mouse reference genome GRCm38.102 (mm10). General differential expression analysis was using DESeq2 ^64^ and DEBrowser ^65^ (ViaFoundry). The differential expression analysis was after filtering out genes with counts <1 in majority of samples retaining at least 10,000 genes, followed by Combat batch effect correction, as enabled within DEBrowser. The A>G edits were identified in the merged bam files, processed using the SAILOR pipeline ^29,33^, with following input parameters: “junction_overhang“: 10, “edge_mutation“: 5, “non_ag“: 0, “dp“: “DP4”, “min_variant_coverage“: 5, “edit_type“: “ag”, “edit_fraction“: 0.01, to quantify the A to G changes. The SAILOR pipeline results in ascending confidence marked by a score of 0 to 1, though the score cutoff used for individual analysis is mentioned within the results, in most cases only sites with score ≥0.99 are considered. Annotation of the edit sites, also referred to as “peaks” or “single-nucleotide peaks”, was done by ChIPseeker ^66^ with “tssRegion = c(0, 0)” and “sameStrand = TRUE”, as the analysis was for RNA peaks, and gene-body metaplots of the peaks were plotted using MetaPlotR ^67^. Search for RBP binding motif was using HOMER ^68^, usually with parameters “-size 100 -len 6”.

Identification and quantification of differential and alternate use of 3’UTR and polyadenylation sites was with APAlyzer ^45^, for the 3’UTR APA the parameters used are “Strandtype=“forward”, CUTreads=10, p_adjust_methods=“BH”, MultiTest=’unpaired t-test’”, and an additional parameter for IPA is “SeqType=“ThreeMostPairEnd“”. General alternative splicing analysis was done using rMATS-turbo ^44^ (4.3.0). Binding motif distribution around the edit site was analyzed and quantified using annotatePeaks.pl from HOMER tools, using the parameters of -size 200 -hist 10.

### Data availability

Raw and processed RNA sequencing data generated in this study have been submitted to the NCBI GEO database and can be accessed under accession number GSE329111.

## SUPPLEMENTAL MATERIAL

**Supplemental Tables S1-S3**

**Supplemental Figures S1-S6**

## SUPPLEMENTAL TABLE LEGENDS

**Table S1. RNA-seq for the EERRi dataset.**

Sheets show DESeq2 output for matched sets of rABE-only and rABE-RBP-injected embryos, with 17 sheets for the RBPs under investigation in this study.

**Table S2. Editing dataset.**

Sheets show edit counts for each of the 107 datasets generated for rABE-injected embryos, with sheets providing data for edits at P>0.9 and P>0.99 confidence levels.

**Table S3. RNA-seq for RBP knockdowns.**

DESeq2 output for scramble and anti-RBP knockdown experiments.

## SUPPLEMENTAL FIGURE LEGENDS

**Figure S1.**
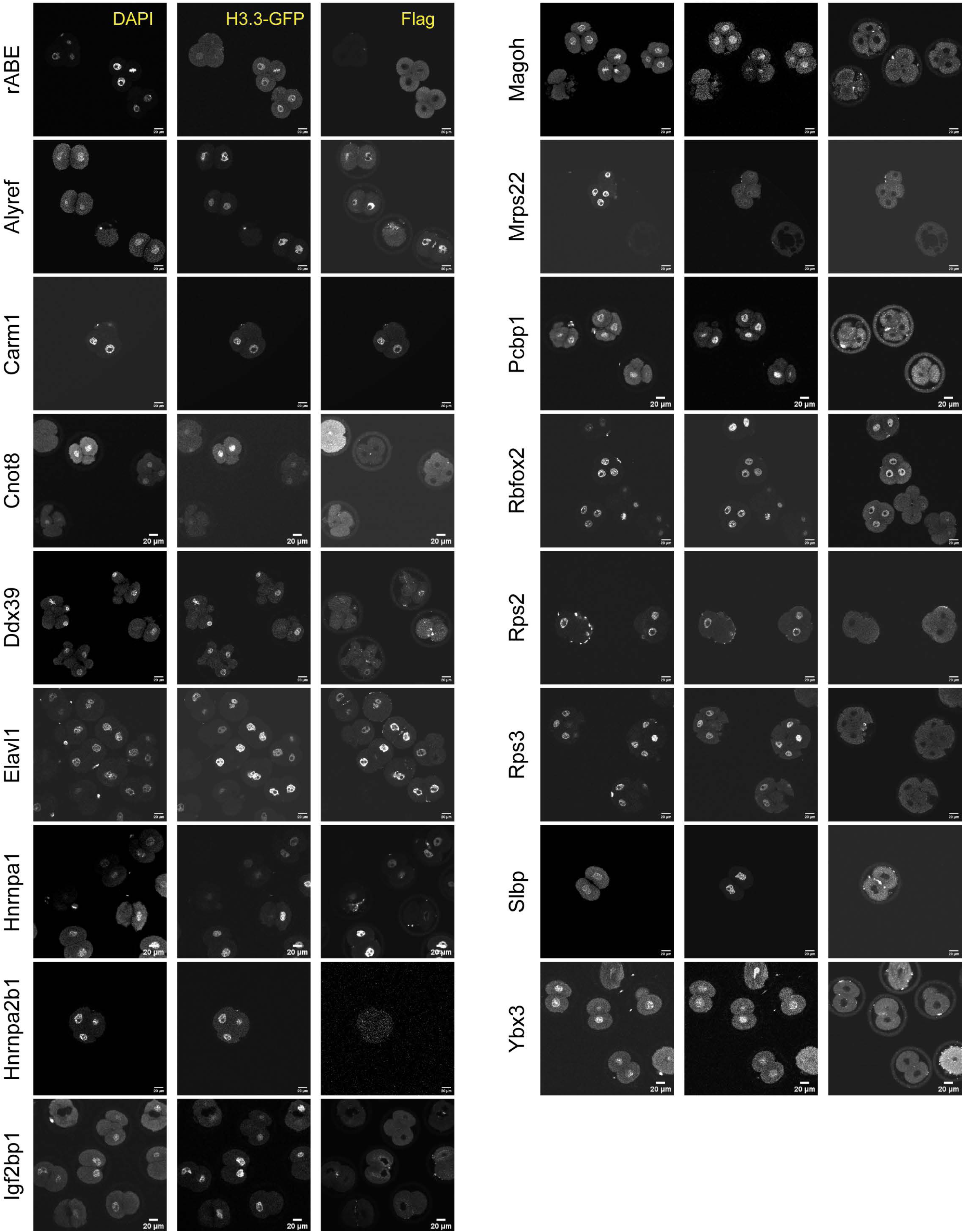
Subcellular localization of RBP-rABE fusion proteins. For each RBP, 3 panels show DAPI staining, H3.3-GFP (to visualize nuclei), and RBP-rABE (tagged with a C-terminal Flag epitope).

**Figure S2.**
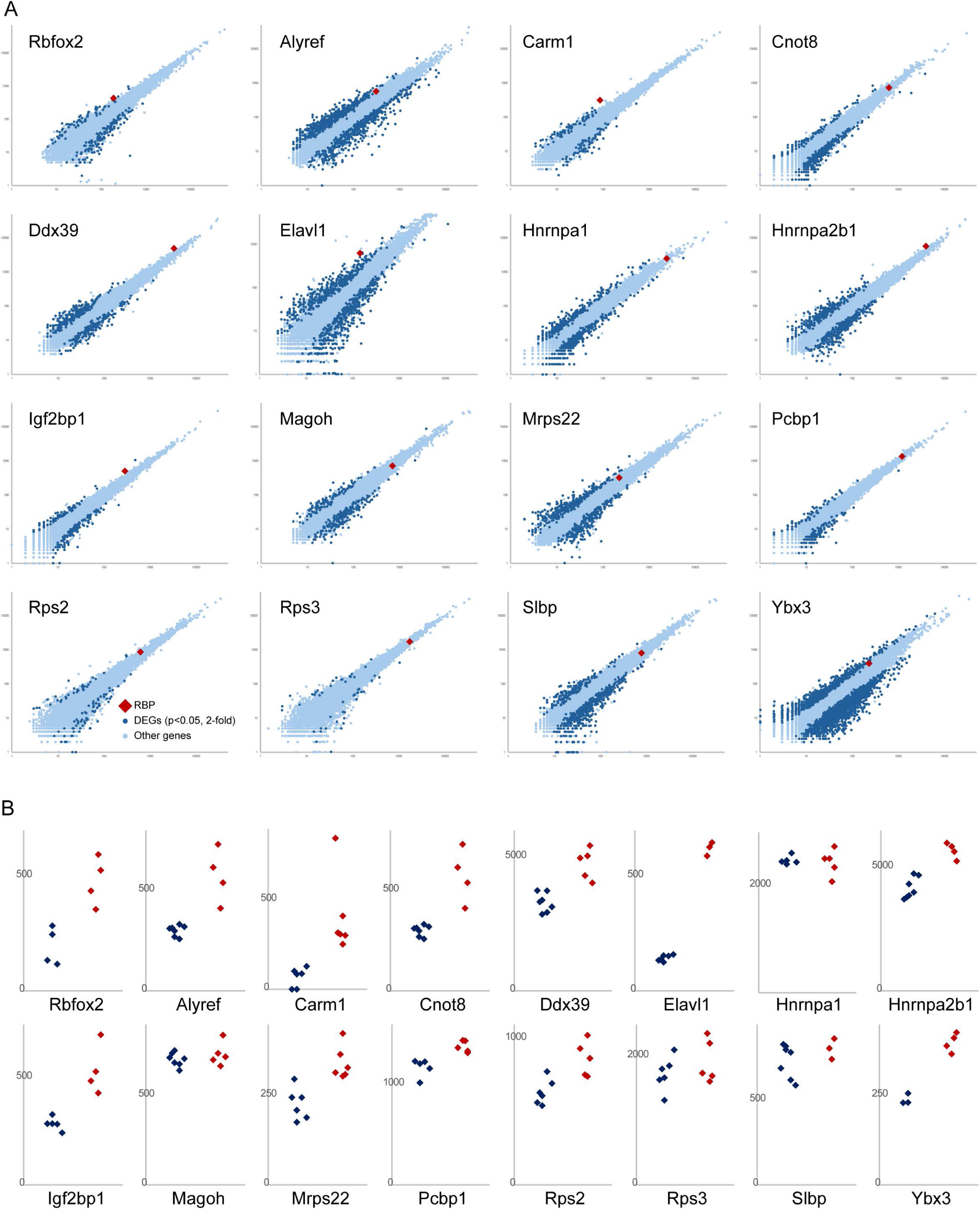
Effects of RBP-rABE expression on embryo gene regulation. A) Scatterplots show mRNA abundance comparing rABE-only expressing embryos (x axis) with RBP-rABE-injected embryos (y axis). In each case the mRNA encoding he RBP of interest is highlighted in red. B) Dot plots show expression levels (tpm) for each RBP for RBP-rABE-injected embryos (red dots) and matched rABE-only controls (blue dots).

**Figure S3.**
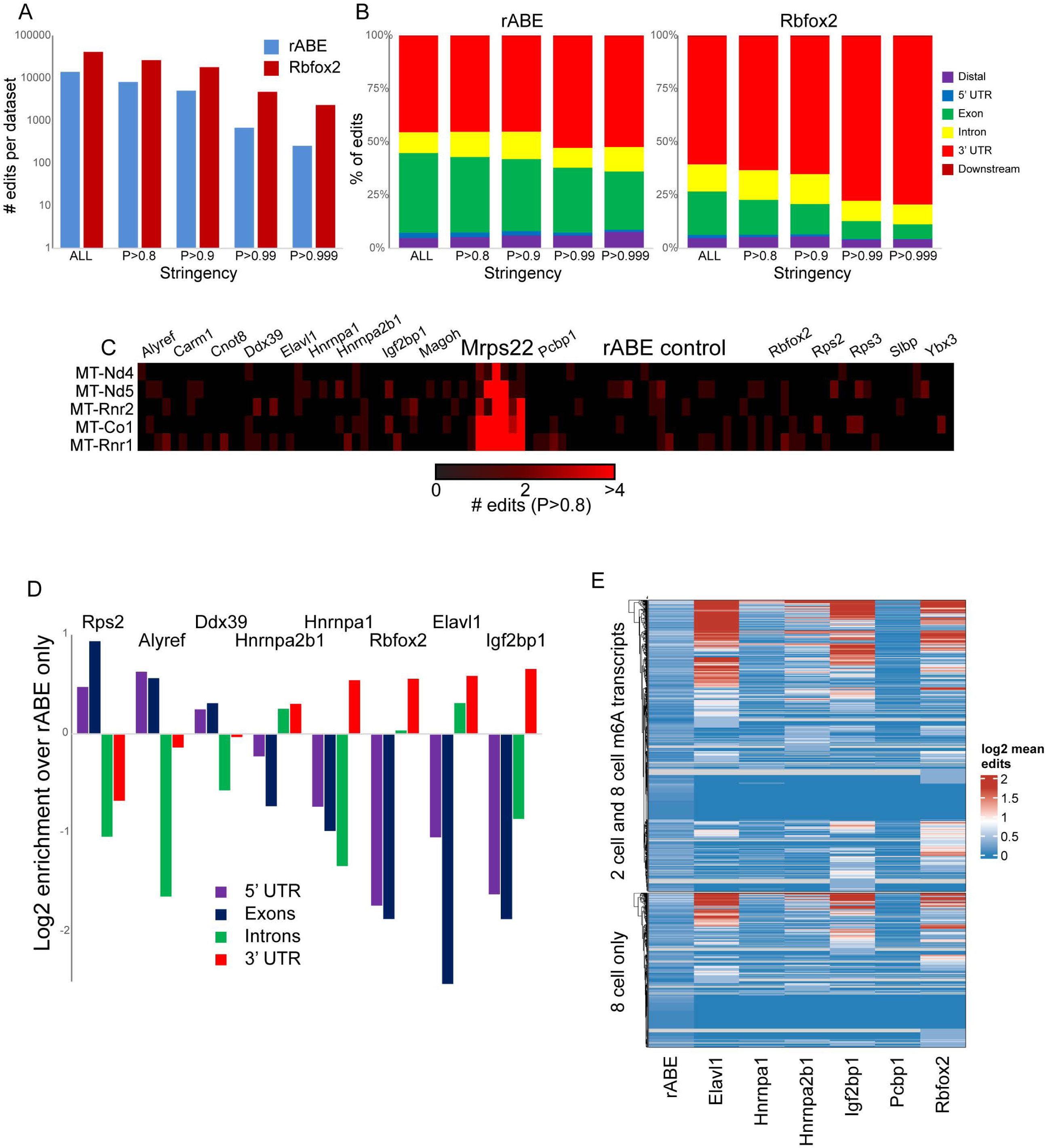
Expected features of EERRi datasets. A) Average number of edits per dataset for Rbfox2-rABE, or rABE-only control, at the indicated stringency for editing calls. B) Localization of edits across gene models for edits of varying stringency. C) Heatmap shows edit counts for all detected mitochondrial transcripts across all datasets, confirming that mitochondrial edits are confined to the Mrps22 dataset. D) Enrichment for RBP edits (calculated as Log2 FC RBP-rABE/rABE only, at an editing stringency of P>0.99) across the indicated genic regions. E) Overlap between m6A-marked transcripts from Wang, Li et al, Table S1 ^37^, and RNA editing for the indicated RBPs. Heatmap shows all transcripts called as m6A-labeled at the 8-cell stage alone, or at both 2-cell and 8-cell stages, and color bar shows the editing ratio (compared to rABE alone) for the indicated RBPs.

**Figure S4.**
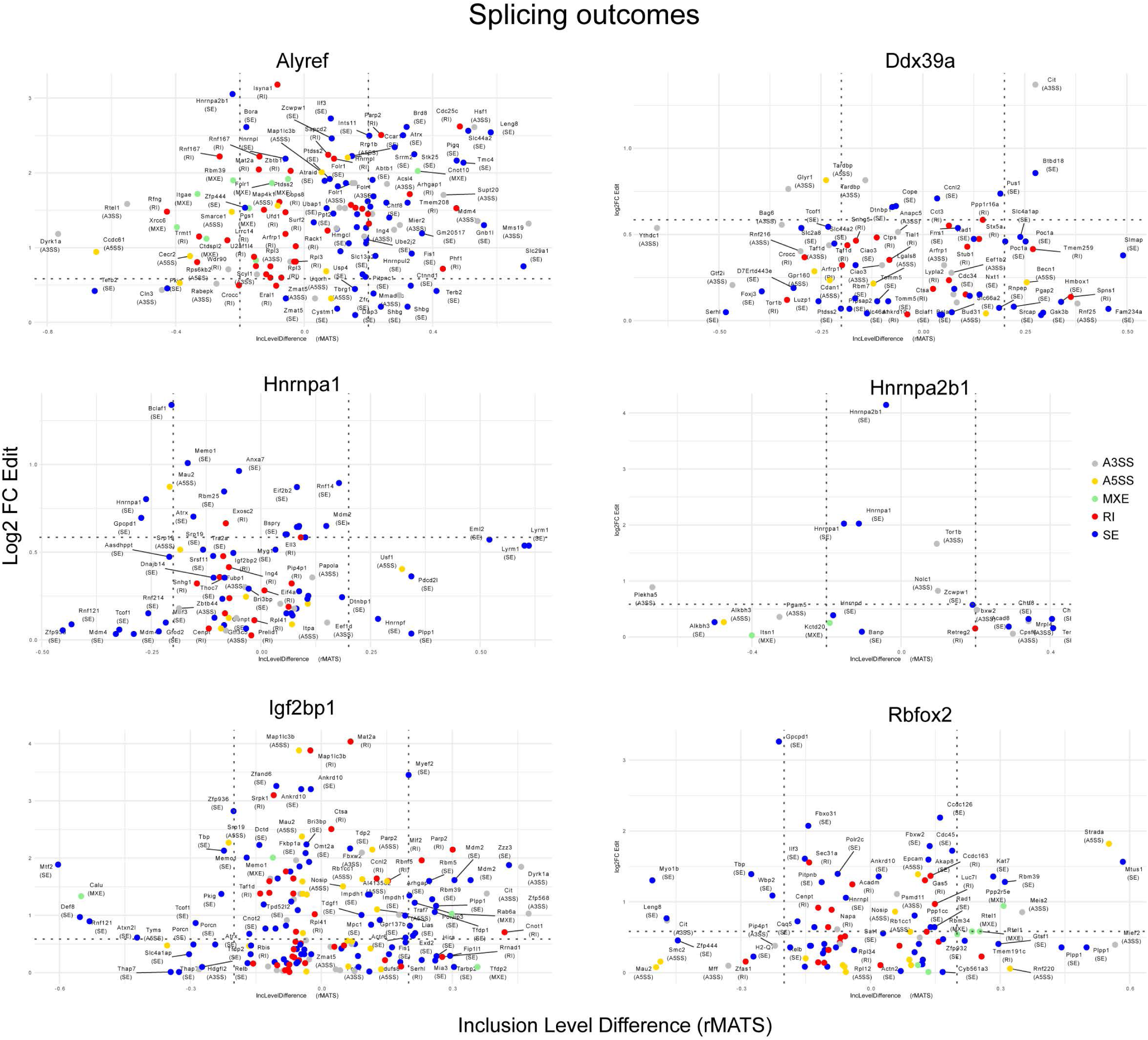
Splicing outcomes for the indicated RBP knockdowns. Scatterplots comparing altered splicing (x axis) vs. RBP-rABE-directed edits (y axis), as in Fig. 5B.

**Figure S5.**
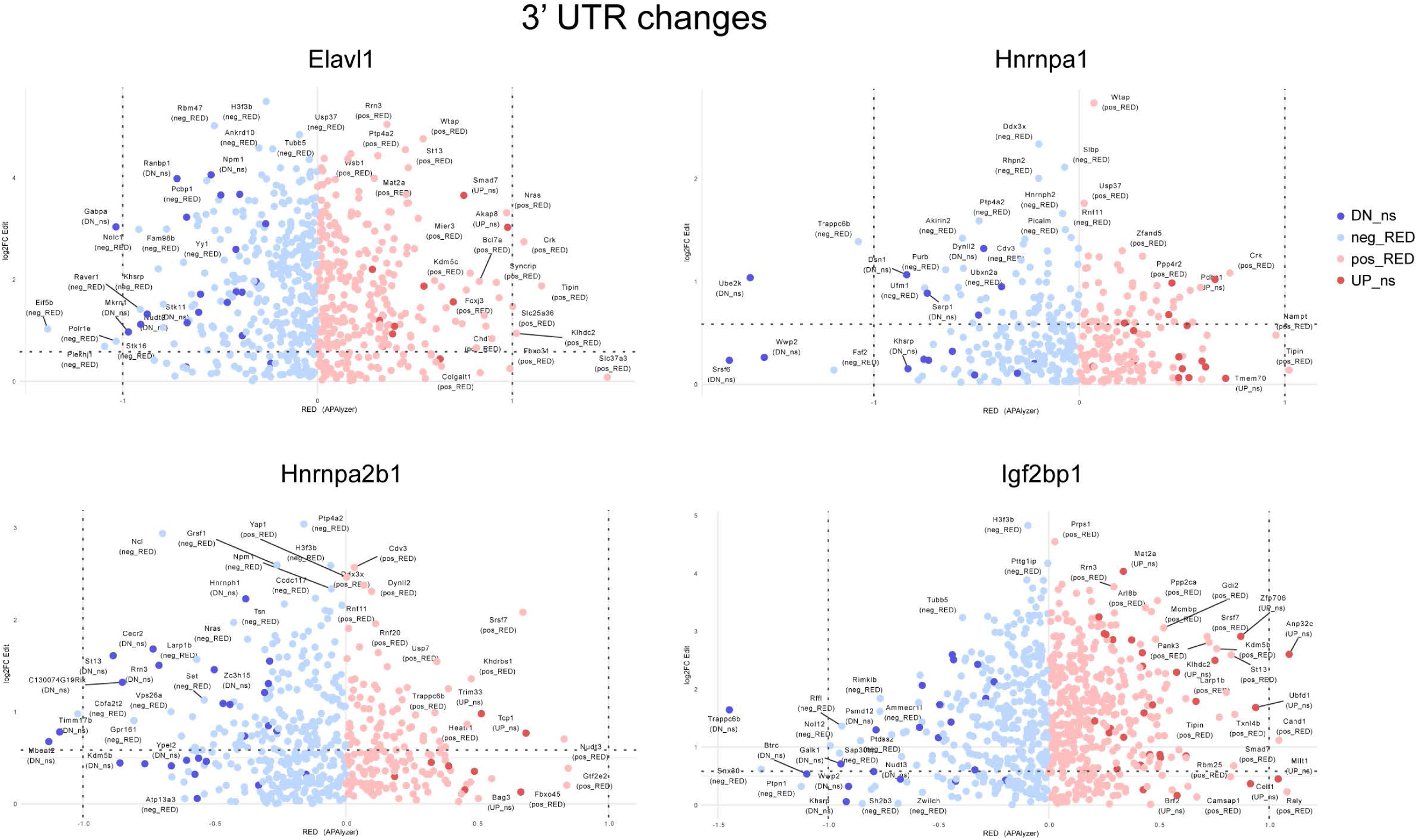
Alternative polyadenylation outcomes following RBP knockdowns. Changes in 3’ UTR length are plotted for the indicated RBP knockdowns, as in Fig. 6A.

**Figure S6.**
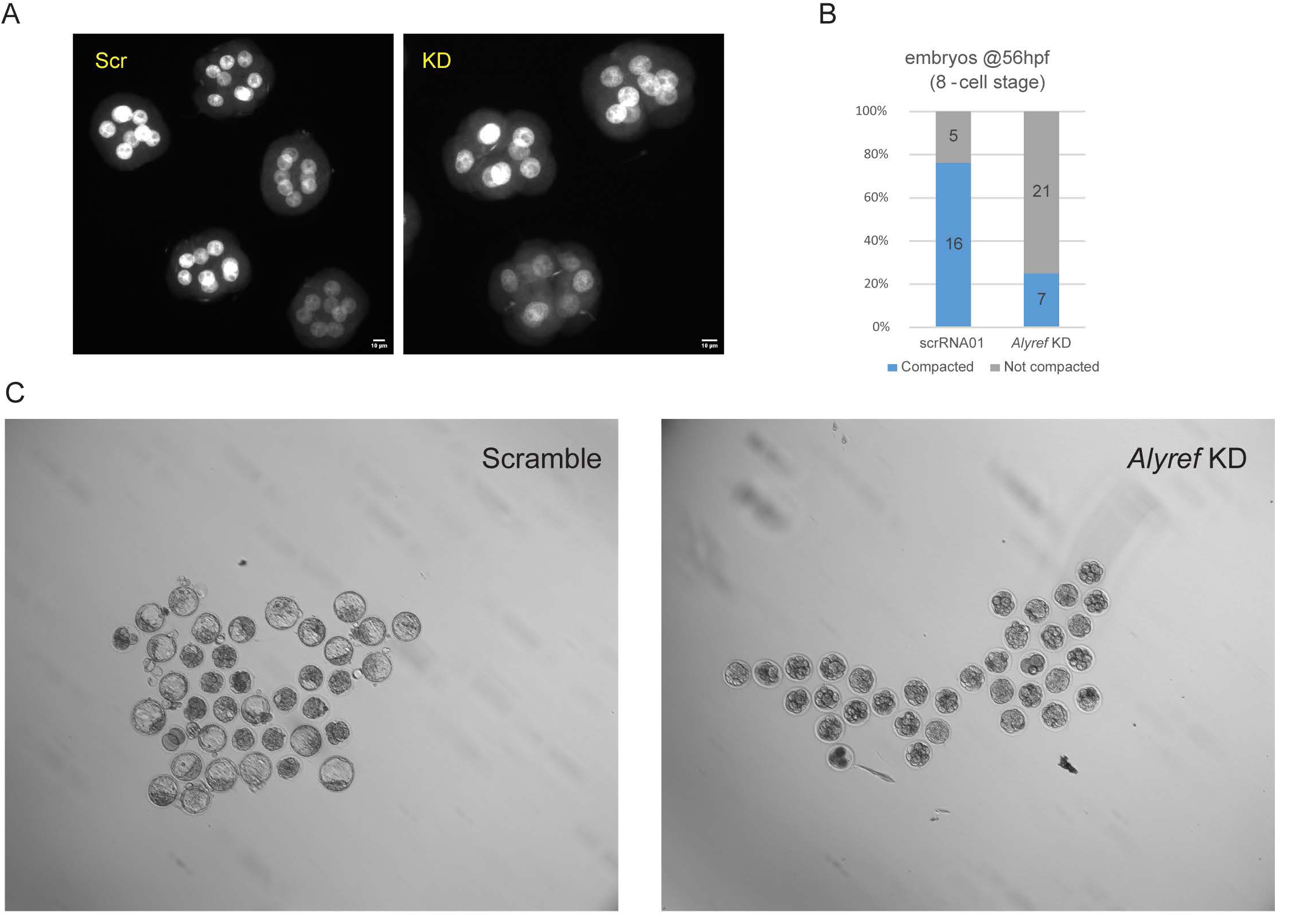
Alyref is required for normal Ddx21 expression and preimplantation development. A) *Alyref* KDs exhibit poor compaction at the eight cell stage. DAPI-stained images show typical control and KD embryos. B) Quantitation of embryo compaction at 56 hpf. C) *Alyref* is required for blastocyst progression, here visualized as blastocyst cavity formation. Images are for embryos 96 hpf.

